# wenda_gpu: fast domain adaptation for genomic data

**DOI:** 10.1101/2022.04.09.487671

**Authors:** Ariel A. Hippen, Jake Crawford, Jacob R. Gardner, Casey S. Greene

## Abstract

**Motivation:** Domain adaptation allows for development of predictive models even in cases with limited sample data. Weighted elastic net domain adaptation specifically leverages features of genomic data to maximize transferability but the method is too computationally demanding to apply to many genome-sized datasets.

**Results:** We developed wenda_gpu, which uses GPyTorch to train models on genomic data within hours on a single GPU-enabled machine. We show that wenda_gpu returns comparable results to the original wenda implementation, and that it can be used for improved prediction of cancer mutation status on small sample sizes than regular elastic net.

**Availability:** wenda_gpu is available on GitHub at https://github.com/greenelab/wenda_gpu/.

**Supplementary information:** Supplementary data are available at *Bioinformatics* online.

## 1 Introduction

Prediction models generated from genomic data are widely used in bioinformatics, including calling mutation status, identifying disease subtype, and predicting cancer prognosis. A fundamental assumption in supervised machine learning is that the data to be classified is derived from the same distribution as the data used to train the classifier. Prediction in a dataset that does not meet this assumption is prone to problematic extrapolation. However, challenges in acquisition of biological data often mean few or no labeled examples are available for the distribution of interest. For instance, sample sizes may be insufficient to train on rare cancer types, or technological limitations may hinder label generation, for instance a lack of simultaneous profiling of gene expression and mutation information in single-cell data. For such situations where labeled target data is limited, the field of domain adaptation and transfer learning has established principled ways to develop predictors for the data of interest (target data) using labeled data from a similar but distinct distribution (source data).

One recent method, weighted elastic net domain adaptation or wenda (Handl *et al*., 2019), leverages the complex interactions between biological features (such as genes) to optimize a model’s predictive power on both source and target datasets. It learns the dependency structure between features and prioritizes those that are similar across source and target distributions. This has previously been shown to significantly improve accuracy on predictions from a mismatched distribution, overcoming the limitations of traditional supervised models. Unfortunately, wenda requires training a Gaussian process model for each feature separately, which is computationally expensive and resists parallelization, making it infeasible for researchers to use at genome-scale.

To address this limitation, we implemented a modified form of the underlying algorithm, called wenda_gpu, which allows for fast, efficient model training for genome-scale datasets on a single GPU-enabled computer.

## 2 Results

### 2.1 Implementing wenda with gpytorch

Weighted elastic net domain adaptation has three major steps. First, each feature *f* in the source distribution and generates a Gaussian process model to predict the expression level of that gene using all other genes (¬*f*). By using a Bayesian model, wenda returns a probability distribution of values © The Author 2015. *P* (*f*_*source*_|¬*f*_*source*_). Next, wenda uses this model to calculate the probability of each sample’s observed value of *f* in the target distribution, *P* (*f*_*target*_|¬*f*_*target*_). A high probability indicates that the relationship between *f* and other key genes is similar between source and target distributions and suggests that *f* will be useful for prediction in the target distribution. Finally, these probabilities are averaged over all target distribution samples and transformed into a weight for each gene, which is included in the penalization term for the ultimate elastic net predictor (Handl *et al*., 2019). This first step, generating a Gaussian process model for each feature, is highly time-intensive. We implemented the model training step using GPyTorch, which provides efficient and modular Gaussian process inference (Gardner *et al*., 2021).

### 2.2 wenda_gpu implementation reduces runtimes

To assess the time cost of native wenda, we ran on the data used in the original paper predicting age from methylation data on numerous tissues. The authors of the original paper reported that training models for 12980 methylation sites across 1866 samples took 51 hours on 10 CPUs (Handl *et al*., 2019). When we ran native wenda on a computer with 6 CPUs, we allowed the program to run for 3 days, during which it trained 1309/12980 models before becoming unresponsive. Alternatively, when we ran wenda_gpu on the same computer, equipped with a Titan Xp GPU, all models trained in approximately 5 hours. To more precisely quantify the speed increase, we ran native wenda and wenda_gpu on several smaller datasets of various sizes (Fig. 1A). For instance, on small datasets of 500 features, all models trained in 836 seconds on native wenda and 116 seconds on wenda_gpu (average across three replicates), meaning an approximately 7-fold decrease in time to run. On a slightly larger dataset with 2000 features, native wenda trained all models in 6851 seconds as opposed to 554 seconds on wenda_gpu, a 12-fold speedup. Using wenda_gpu, completing the whole prediction task on genome-wide datasets with tens of thousands of features is thus feasible in a single day on a single GPU-enabled computer.

**Fig. 1.**
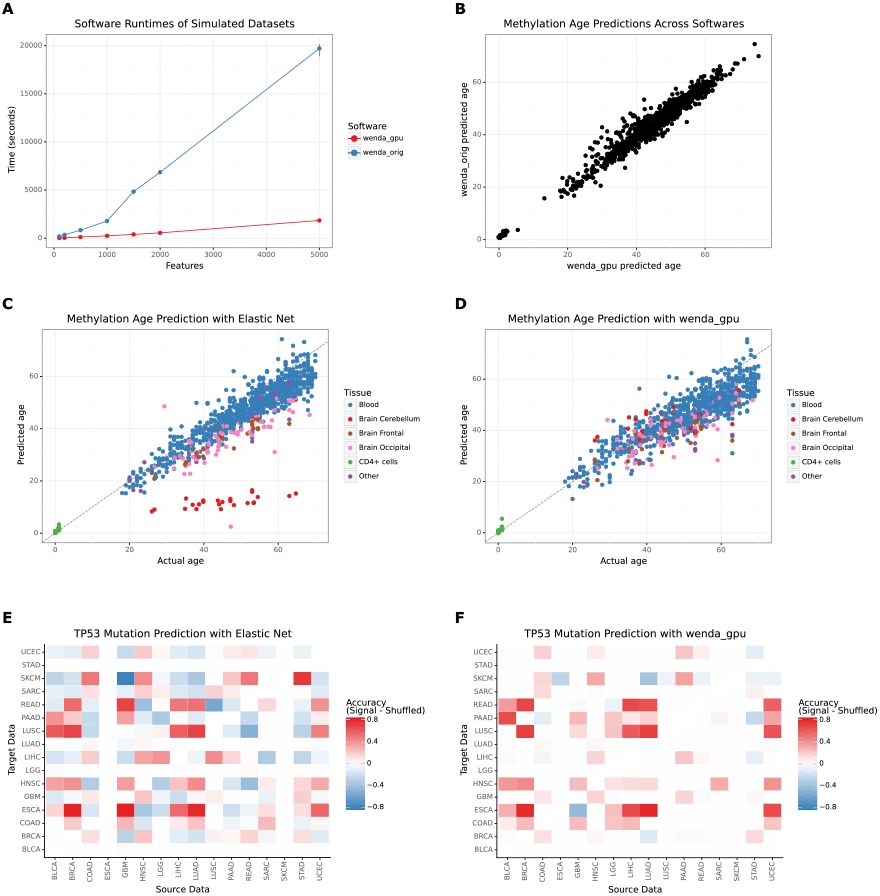
(A) Mean runtimes in wenda_orig and wenda_gpu to train feature models and generate confidence scores for simulated datasets (3 replicates for each number of features). (B) Correlation between wenda_gpu and wenda_orig on methylation age prediction at k=6. X axis is one of 10 cross-validation folds from wenda_orig, chosen for being closest to median correlation value of 0.977. (C, D) Age predictions on methylation data done by (C) regular elastic net and (D) wenda_gpu, compared with actual age. With a regular elastic net, prediction is poor on cerebellum tissue (red), which is not in the source dataset. Using wenda_gpu (k=6), cerebellum samples are predicted well, even though the model was not provided with labels for this tissue type. (E, F) Difference in accuracy of TP53 mutation prediction on pairs of TCGA cancer types compared to a model with shuffled labels, done using elastic net (E) and wenda_gpu (F).

### 2.3 wenda_gpu replicates original wenda results

To confirm that wenda_gpu returned similar results for the ultimate classification task, we compared results from the elastic net classifiers from wenda_gpu and native wenda. The age predictions between the two methods were highly correlated (median r=0.977 for k=6) (Fig. 1B). Consistent with the original wenda manuscript, we found that a classifier trained with weighted elastic net domain adaptation was more accurate on a tissue type not found in the source data than was a regular elastic net (Fig. 1C).

### 2.4 wenda_gpu enables domain adaptation in new contexts

With the improved speed of wenda_gpu, we were able to assess the utility of weighted elastic net domain adaptation in new contexts. One example use case is building classifiers on transcriptomic data to identify mutations in common cancer driver genes, which has been shown to be possible in a pan-cancer context (Way *et al*., 2018; Knijnenburg *et al*., 2018). Mutation classifiers trained on transcriptomic data from one cancer type can even be predictive of mutation status in other cancer types (Crawford *et al*., 2021). We postulated that wenda would allow prediction models trained on one cancer type to perform better on another cancer type, opening the possibility of these models being used on rare cancers for which sample sizes are limited. To explore this, we trained classifiers to identify mutations in TP53, the most commonly mutated gene in cancer (Mendiratta *et al*., 2021). We selected 16 cancer types from TCGA with sample size greater than 100 and with between 10% and 90% of samples having a TP53 loss-of-function mutation. We trained models pairwise (e.g. for each two cancer types A and B, train a model using transcription data from A as source data and B as target data, and vice versa) using wenda_gpu and a regular elastic net. We found that the models trained with wenda_gpu had equal or higher accuracy than the elastic net models in 201/240 pairs, with a pronounced improvement in some cancer type pairs. As a negative control, we also trained the same models with shuffled TP53 mutation labels. Across all cancer type pairs, models trained with wenda_gpu more consistently outperformed the negative control than did the regular elastic net models (Fig. 1D).

## 3 Conclusion

Domain adaptation allows researchers to leverage areas with abundant data to enable machine learning methods in use cases with more limited data. Weighted elastic net domain adaptation exploits the complex biological interactions that exist between genomic features to maximize transferability to a new context. With the increase in speed afforded by wenda_gpu, researchers can apply wenda to larger datasets and experiment with more potential use cases of domain adaptation on genomic data without a significant expense of researcher time. The wenda_gpu software is available at https://github.com/greenelab/wenda_gpu/. The code used to perform all analyses in this paper is available at https://github.com/greenelab/wenda_gpu_paper/.

## Acknowledgements

The authors would like to thank Lisa Eisenberg for consulting on a new implementation of wenda, as well as providing methylation data and *wenda_gpu: fast domain adaptation for genomic data* **3** intermediate model confidence scores. Thanks also to Ben Heil, Alexandra Lee, and David Nicholson for code review.

## Funding

AAH and CSG and were supported by the National Cancer Institute (NCI) from the National Institutes of Health (NIH) grant R01CA237170. JC and CSG were also supported by grant R01HG010067 from the NIH’s National Human Genome Research Institute (NHGRI). The funders had no role in study design, data collection and analysis, decision to publish, or preparation of the manuscript.

## Notes

https://github.com/greenelab/wenda_gpu/

### Competing Interest Statement

The authors have declared no competing interest.

